# Targeting aberrant dendritic integration to treat cognitive comorbidities of epilepsy

**DOI:** 10.1101/2020.11.23.393694

**Authors:** Nicola Masala, Martin Pofahl, André N Haubrich, Khondker Ushna Sameen Islam, Negar Nikbakht, Maryam Pasdarnavab, Fateme Kamali, Kirsten Bohmbach, Christian Henneberger, Kurtulus Golcuk, Laura A Ewell, Sandra Blaess, Tony Kelly, Heinz Beck

## Abstract

Memory deficits are a debilitating symptom of epilepsy, but little is known about mechanisms underlying cognitive deficits. Here, we describe a Na^+^ channel-dependent mechanism underlying altered hippocampal dendritic integration, degraded place coding, and deficits in spatial memory.

Two-photon glutamate uncaging experiments revealed that the mechanisms constraining the generation of Na^+^ spikes in hippocampal 1^st^ order pyramidal cell dendrites are profoundly degraded in experimental epilepsy. A selective Na_v_1.3 sodium channel blocker reversed this effect, and Nav1.3 channels were up-regulated selectively in principal neurons at the mRNA level. Finally, in-vivo two-photon imaging revealed that the Na_v_1.3 channel blocker improves degraded hippocampal spatial representations, and reverses hippocampal memory deficits.

Thus, a dendritic channelopathy may underlie cognitive deficits in epilepsy and targeting it pharmacologically may constitute a new avenue to enhance cognition.

**One Sentence Summary:** Impaired input computations via aberrant dendritic spikes in chronic epilepsy degrade neuronal place codes and spatial memory

## Main Text

In most CNS pyramidal neurons, dendrites are capable of generating spikes in local membrane potential that are initiated by dendritic voltage gated Na^+^ channels (*1*). Because dendritic spikes are elicited by spatiotemporally clustered inputs, arising only if specific ensembles of presynaptic neurons are synchronously active, they have been proposed to endow neurons with the capability to act as input feature detectors (*2*). Indeed, dendritic spikes have been found to be relevant for triggering place-related firing in CA1 neurons (*3, 4*). They thus constitute a key mechanism for dendritic integration and neuronal input-output computations, and have been strongly implicated in spatial navigation.

Dendritic spikes rely on targeted expression of voltage-gated ion channels in dendritic branches. In epilepsy, as well as in numerous other CNS disorders, the expression and function of ion channels are profoundly altered in CA1 neuron dendrites. In chronic epilepsy models, changes in K^+^ channels (*5*), T-type Ca^2+^ channels (*6, 7*), HCN channels (*8, 9*) and Na^+^ channels have been identified (*10*). However, these studies have mainly focused on larger caliber, apical dendrites of pyramidal neurons, primarily because of the difficulties in obtaining direct patch-clamp recordings from thin dendrites. The integrative properties of thin, higher-order dendrites, and how they change in chronic epilepsy have not been studied so far.

In this study, we address how dendritic integration via dendritic spikes is altered in chronic epilepsy, how this affects neuronal coding in vivo, and how it impacts behavior. We propose that an up-regulation of Na^+^ channels in hippocampal pyramidal neuron apical oblique dendrites underlies altered dendritic spikes, degraded place coding, and cognitive deficits in experimental temporal lobe epilepsy (TLE).

## Results

### Altered dendritic integration via dendritic spikes in chronic epilepsy

We examined dendritic integration in the kainate model, a well-established model of chronic temporal lobe epilepsy (*11–13*), characterized by spontaneous recurrent seizures (**Fig. S1a-c**) and hippocampal pathology (**Fig. S2**). The intrinsic firing properties of neurons showed an increased action potential output (**Fig. S3a-c**), as well as an increased fraction of bursting neurons, similar to other mouse TLE models (*6*) (**Fig. S3d,e, Table S1**).

To probe dendritic integration in CA1 pyramidal neurons we used two-photon glutamate uncaging (*1, 14*). Responses of single spines to uncaging of glutamate (uEPSPs), were calibrated to ~1 mV in both sham-control and epileptic animals (1.09±0.42 mV, n=98 vs. 1.07±0.04 mV, n=85 dendrites, unpaired t-test, *p*=0.80, **Fig. S4a-c**). The rise and decay kinetics of such single-spine unitary EPSPs elicited by uncaging were slightly but significantly faster in epileptic animals (**Fig. S4d, e**). This was also observed in miniature EPSC (mEPSC) recordings (**Fig. S4f-k**). Spine density in 1^st^ and 2^nd^ order dendrites did not differ between sham-controls and epileptic animals (**Fig.S4l-n**).

We then went on to probe the capability of CA1 dendrites to generate dendritic spikes by stimulating multiple spines quasi-synchronously (interspine stimulation interval, 0.1ms). As described previously (*1, 14*), CA1 dendrites either displayed linear integration only, or were capable of generating sudden supralinear dendritic spikes when increasing numbers of spines were synchronously stimulated (representative examples in **Fig. 1a-c**). In linearly integrating dendritic branches, the linear summation of single spine uEPSPs was augmented in epileptic animals in 1^st^ order branches emanating from the main dendritic shaft (n=35 and 26 in sham-control vs. epileptic mice). Linearly integrating 2^nd^ order dendrites did not show differences in EPSP summation (**Fig. 1d, f**, n=23 and 15 in sham-control vs. epileptic mice, two-way ANOVA main effect, sham-control vs. epileptic mice: F_(1 95)_=7.38, *p*=0.0078; 1^st^ order vs. 2^nd^ order: F_(1 95)_=4.33, *p*=0.040; interaction: F_(1 95)_=0.32, *p*=0.57; Bonferroni’s post-test, 1^st^ order dendrites sham-control vs. Epileptic mice *p*=0.018; 2^nd^ order dendrites sham-control vs. epileptic mice *p*=0.35).

**Fig. 1:**
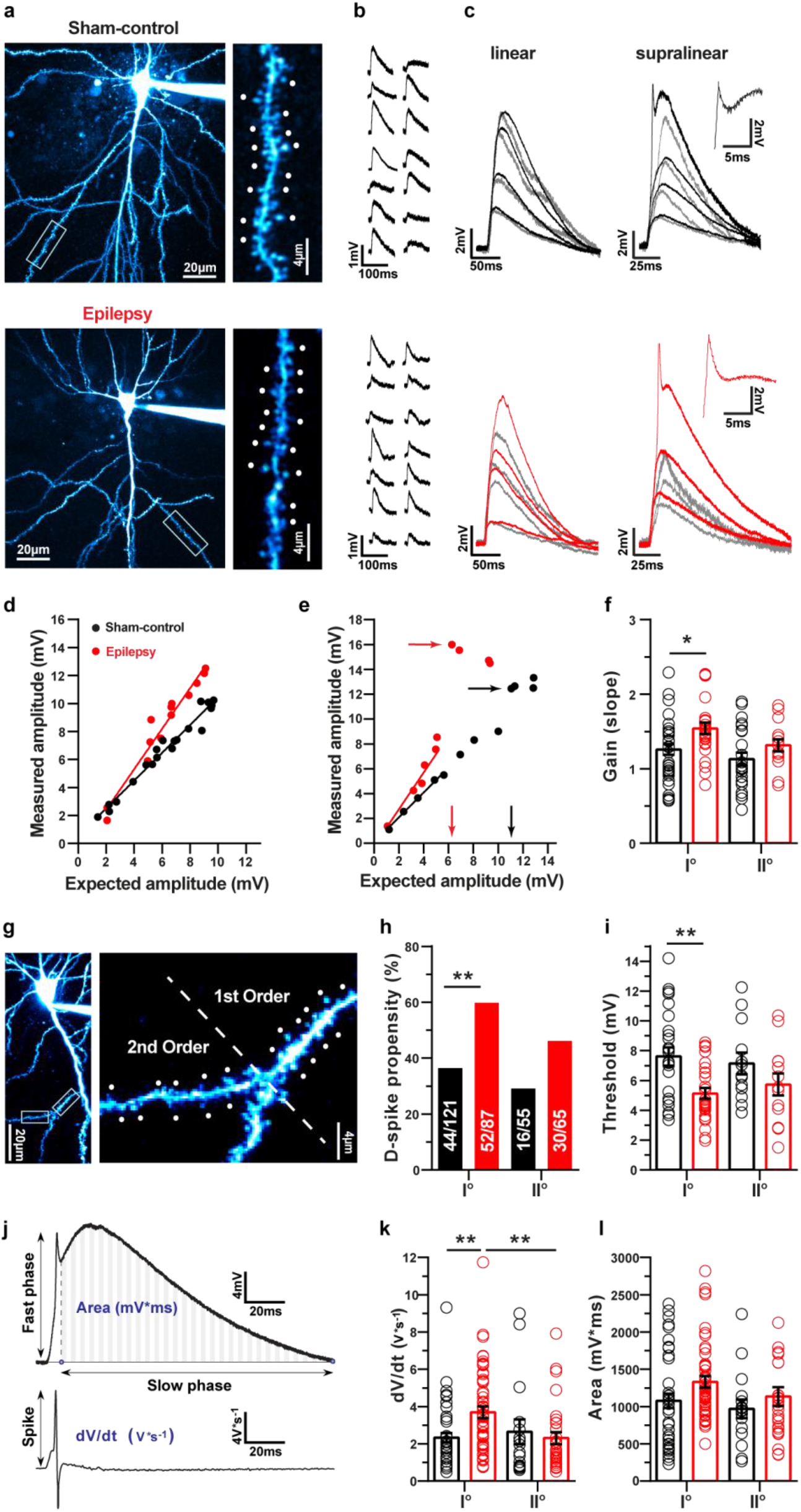
Probing dendritic integration in chronic epilepsy (kainate model of epilepsy). **a,** CA1 pyramidal neurons filled with a fluorescent dye via the somatic patch recording, close up of a dendritic segment with uncaging targeting points at spines. **b,** Responses to single-spine stimulation with 2P-uncaging of MNI-glutamate measured with somatic patch recording. **c,** Representative recordings of compound linear and supralinear EPSPs. Grey lines are expected EPSPs from linear summation of single spine responses. Examples of a branch with linear integration (left panels) and a branch capable of supralinear integration (right panels) are shown for sham-control (black) and epileptic mice (red). **d,** Examples of linearly integrating dendrites in sham-control (black) and epileptic animals (red). **e,** Examples of dendrites capable of generating supralinear dendritic spikes in sham-control (black) and epileptic animals (red). Occurrence of dendritic spikes and their voltage threshold are indicated with arrows. **f,** Quantification of the slope of the linear phase in linearly integrating 1^st^ and 2^nd^ order dendrites in sham-control (black) and epileptic mice (red). Asterisks indicate Bonferroni’s post-tests *p*=0.0176. **g,** Representative dendrite with 1^st^ and 2^nd^ order branches. **h,** The propensity for dendritic spikes is enhanced in 1^st^ order dendrites in epileptic animals (red) compared to sham-controls (black). Asterisks indicate Fisher’s exact test *p*=0.0011. **i,** The threshold for generation of dendritic spikes, measured as indicated with arrows in panel e is reduced in 1^st^ order dendrites of epileptic animals, but not in 2^nd^ order dendrites. Asterisks indicate Bonferroni’s post-test, *p*=0.0017. **j-l,** The fast phase of dendritic spikes is accelerated in 1^st^ order dendrites from epileptic mice (panel k, asterisks indicate Bonferroni’s post-test, rate of rise in 1^st^ order dendrites in sham-control vs. epileptic *p*=0.0025; 1^st^ vs. 2^nd^ order dendrites rate of rise in epileptic *p*=0.0074). The area of the slow phase was unchanged.

The difference in linear summation was normalized by application of the Na^+^ channel blocker tetrodotoxin (TTX), indicating that it is caused by increases in voltage-gated Na^+^ currents (**Fig. S5**, sham-control and epileptic n=7 and 6, respectively, two-way repeated measures ANOVA main effect, sham-control vs. epileptic: F_(1 11)_=0.88, *p*=0.37; ACSF vs. TTX: F_(1 11)_=26.96, *p*=0.0003; interaction: F_(1 11)_=10.15, *p*=0.0087; Bonferroni’s post-test, ACSF vs. TTX in epileptic *p*=0.0003).

Strikingly, the fraction of dendrites capable of generating dendritic spikes was significantly increased in 1^st^ order dendrites, but not in 2^nd^ order dendrites in epileptic animals (**Fig. 1e, g, h**, 1^st^ order dendrites 36% vs. 60%, Fisher’s exact test *p*=0.0011, 2^nd^ order dendrites 29% vs. 46%, Fisher’s exact test *p*=0.062). In these experiments, the average distances of the uncaging sites from the somatic region were not different when comparing sham-control and epileptic mice (1^st^ order dendrites sham-control 66.2±2.2 μm, n=101 vs. epileptic mice 64.9±2.3 μm, n=69; 2^nd^ order dendrites sham-control 77.5±2.6 μm, n=54 vs. epileptic mice 80.6±2.8 μm, n=55; unpaired Student’s t-test 1^st^ order sham-control vs. epileptic mice *p*=0.68; 2^nd^ order sham-control vs. epileptic mice *p*=0.43).

We next compared the properties of dendritic spikes in those branches capable of generating them. The threshold for generation of dendritic spikes was computed by determining the expected somatic voltage at which a dendritic supralinearity first occurred (indicated by arrows at the x-axis, **Fig. 1e**). In epileptic animals, the voltage threshold to generate dendritic spikes was significantly reduced in 1^st^ order, but not 2^nd^ order dendrites (**Fig. 1i**, 1^st^ order dendrites n=25 and 27, 2^nd^ order dendrites n=13 and 13 for sham-control and epileptic mice respectively, two-way ANOVA main effect, sham-control vs. epileptic: F_(1 74)_=9.92, *p*=0.0024; 1^st^ order vs. 2^nd^ order: F_(1 74)_=0.012, *p*=0.91; interaction: F_(1 74)_=0.77, *p*=0.38; Bonferroni’s post-test, threshold in 1^st^ order dendrites in sham-control vs. epileptic *p*=0.0017).

The reduction was remarkably pronounced in some 1^st^ order dendrites from epileptic animals, with dendritic spikes sometimes being generated with stimulation of as few as 1-3 spines, and somatic voltages of as little as ~3 mV (see **Fig. S6** for dendritic spike elicited with a single spine stimulation).

A further characteristic of dendritic spikes is the rate of rise of the initial fast phase, that reflects the contribution of voltage-gated sodium channels. The maximal rate of rise was determined from the first derivation of the voltage trace (indicated in **Fig. 1j**). In 1^st^ order dendrites, but not 2^nd^ order dendrites, we observed a significant increase in the maximal rate of rise in epileptic animals (**Fig. 1k**, 1^st^ order dendrites n=44 and 52, 2^nd^ order dendrites n=16 and 30 in sham-control and KA, respectively, two-way ANOVA main effect, sham-control vs. epileptic: F_(1 139)_=1.86, *p*=0.18; 1^st^ order vs. 2^nd^ order: F_(1 139)_=1.96, *p*=0.16; interaction: F_(1 139)_=5.08, *p*=0.025; Bonferroni’s post-test, rate of rise in 1^st^ order dendrites in sham-control vs. epileptic *p*=0.0025; 1^st^ vs. 2^nd^ order dendrites rate of rise in epileptic *p*=0.0074). In contrast to the fast phase of the dendritic spikes, the subsequent slower depolarization was not different in sham-control vs. epileptic animals (**Fig. 1l**, 1^st^ order dendrites n=42 and 47, 2^nd^ order dendrites n=16 and 29, in sham-control and epileptic, respectively, area under the curve of slow depolarization two-way ANOVA main effect, sham-control vs. epileptic: F_(1 130)_=3.51, *p*=0.063; 1^st^ order vs. 2^nd^ order: F_(1 130)_=1.83, *p*=0.18; interaction: F_(1 130)_=0.17). These results collectively show a dramatically augmented excitability of proximal, 1^st^ order dendrites in epileptic animals, reflected in the prevalence and properties of dendritic spikes.

### Reduced synchrony requirement for dendritic spike generation

Synchronous stimulation is required for generation of dendritic spikes in normal pyramidal neurons, and is thought to be critical to detect the coincident activity of specific presynaptic ensembles (*2*). We therefore tested how dendritic spike generation depends on input synchrony by varying the inter-spine stimulation interval in control and epileptic animals (representative examples in **Fig. 2a**, sham-control in black, KA in red). Epileptic animals exhibited a remarkable loss of their capability to selectively respond to synchronous inputs via generation of dendritic spikes. While sham-control animals exhibited a steep decrease in dendritic spike generation when stimulation was less synchronous, epileptic animals continued to generate dendritic spikes even with very asynchronous stimulations (**Fig. 2b,** n=12 and 11 in sham-control vs. epileptic mice, Fisher’s exact test, p<0.001 indicated with asterisks, see figure legend).

**Fig. 2:**
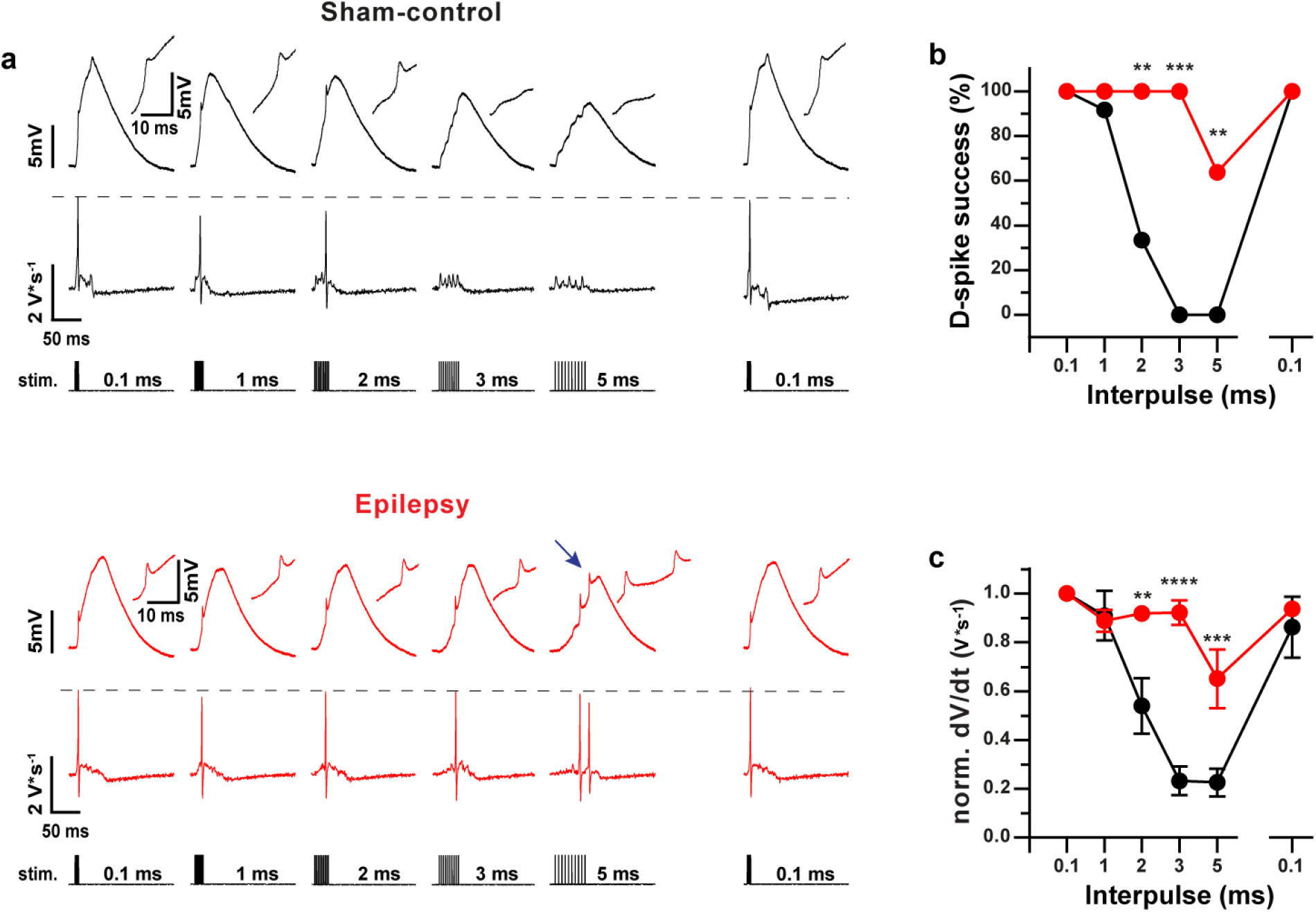
Degraded synchrony requirement in epileptic mice. **a,** Representative example traces showing input synchrony dependence of dendritic spikes in CA1 neurons from sham-control and epileptic mice. Changing the inter-spine stimulation interval (bottom rows in Sham-control and Epileptic) systematically affects dendritic spike generation in control and epileptic animals. Upper rows show somatic voltage response, middle rows show first derivation of the voltage trace (dV/dt). Note that the rightmost response is again elicited with a 0.1 ms inter-spine stimulation interval, applied following the longer intervals. **b, c,** Rate of spike occurrence in percent of uncaging stimulations (D-spike success) and dendritic-spike strength expressed as the maximal rate of rise (dV/dt) of the fast phase of the dendritic spike. Asterisks in b correspond to significance in Fisher’s exact test, with the following p-values: 2 ms *p*=0.0013; 3 ms p<0.0001; 5 ms *p*=0.0013. Asterisks in c correspond to significances in Bonferroni’s post-tests for inter-spine stimulation intervals 2 ms *p*=0.0013; 3 ms p<0.0001; 5 ms *p*=0.0003.

Correspondingly, the maximal rate of rise of the fast phase of dendritic spikes (dV/dt) also showed a much less pronounced reduction with less synchronous stimulation in epileptic animals (**Fig. 2c,** two-way repeated measures ANOVA main effects, sham-control vs. epileptic: F_(1 21)_=32.34, p<0.0001; inter-spine time: F_(4 84)_=22.59, *p*<0.0001; interaction: F_(4 84)_=9.87, p<0.0001; Bonferroni’s post-tests indicated with asterisks, p<0.003, individual p-values see legend). Intriguingly, in some cases, dendritic branches in epileptic animals were capable of generating multiple sequential spikes (n=2 of 11 branches, example in **Fig. 2a,** arrow).

### Reduced dendritic spike inactivation in chronic epilepsy

Sparse dendritic spiking is supported by inactivation of dendritic spike generation by prior activity, a phenomenon that relies on inactivation of dendritic sodium channels. In control animals, dendritic spike inactivation was observed similar to published data (*14*), as indicated by a progressive reduction of dendritic spike dV/dt until dendritic spike failure (black traces in **Fig. S7a,** quantification in **Fig. S7b, c**). In epileptic animals, this reduction in dendritic spike dV/dt as well as inactivation of spike generation was significantly less pronounced (**Fig. S7a-c,** statistical results see figure legend).

Thus, collectively, the results show that dendritic spike generation is strongly enhanced, and that multiple mechanisms that underlie sparse generation of dendritic spikes in normal animals are severely degraded in epilepsy.

### Involvement of Na_v_1.3 channels in augmented dendritic integration

The increased incidence of dendritic spikes, as well as the increased rate of rise of the fast phase of the dendritic spikes suggests up-regulation of Na^+^ channels as a potential mechanism. Na_v_1.3 channels are normally down-regulated in early ontogenesis, but have been shown to exhibit increased expression in epilepsy (*15*). We used the selective Na_v_1.3 blocker ICA-121431 (*16*) (100 nM) to examine the role of this channel subunit in dendritic spikes (**Fig. 3a-h**). In these experiments, we specifically selected uncaging sites on dendrites that exhibited robust and noticeable dendritic spikes to assess if ICA-121431 blocked them. Due to this bias, differences in spike properties between sham-control and epileptic animals are not present in this experiment.

**Fig. 3:**
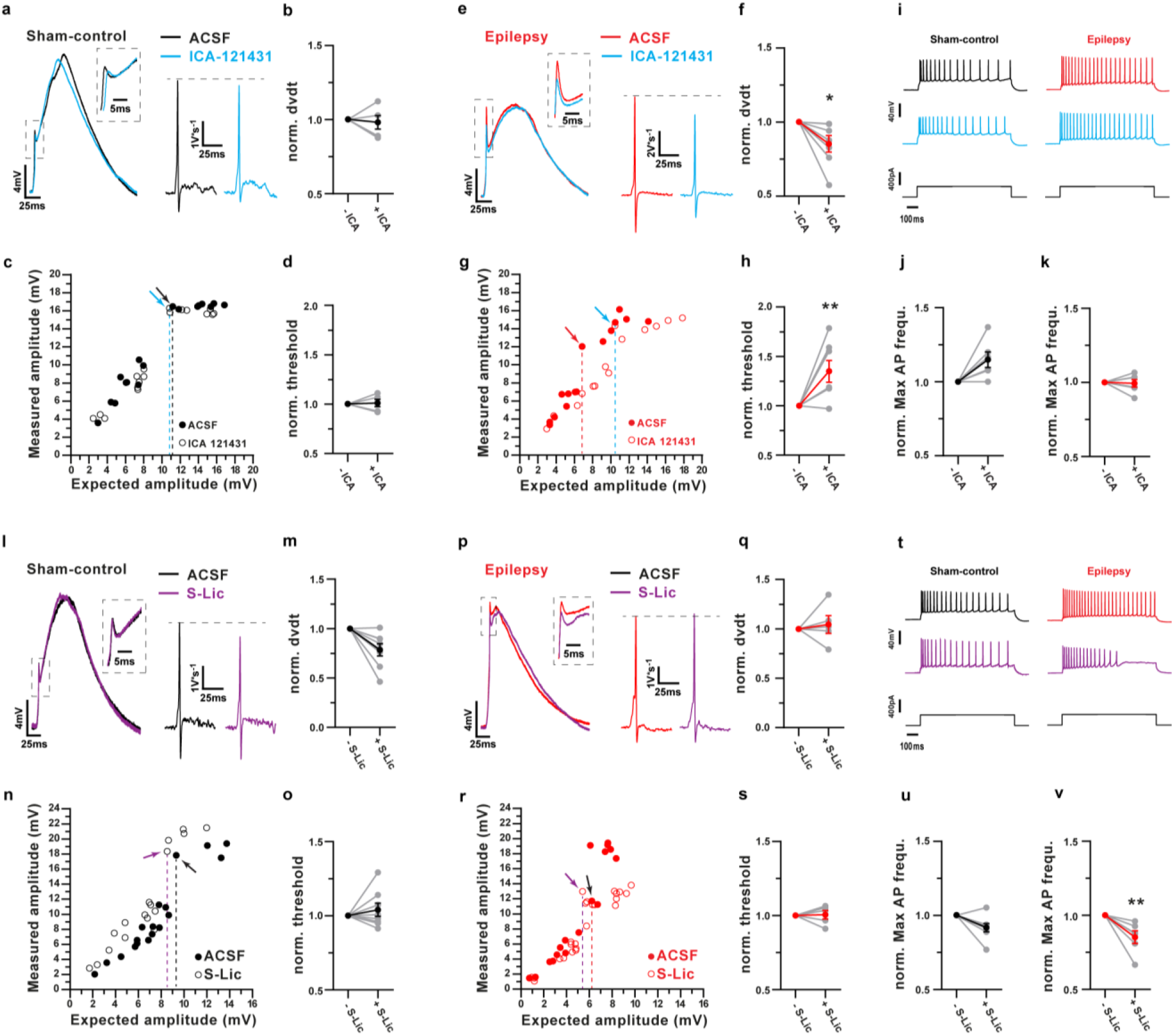
The Na_v_1.3 specific blocker ICA-121431, but not the Na_v_1.2/1.6 Na^+^ channel blocker S-Lic reverses enhanced dendritic excitability. **a, e,** Representative examples of the effects of the Na_v_1.3 specific blocker ICA-121431 (100 nM) on dendritic spikes (insets show magnification of the fast phase of the dendritic spike), and on the first derivation of the voltage trace (dV/dt) in sham-control and epileptic animals. **b, f,** Effects of ICA-121431 at concentrations of 100 nM on the maximal rate of rise of the dendritic spike in sham-control (b) and epileptic mice (f). Asterisk indicates Bonferroni’s post-test, *p*=0.023. **c, g,** Representative examples of effects on the dendritic spike threshold in sham-control and epileptic mice (arrows indicate occurrence of dendritic spikes and dashed lines indicate thresholds). **d, h,** Quantification of the effects of ICA-121431 on the dendritic spike threshold in sham-control (d) and epileptic mice (h). Asterisk indicates Bonferroni’s post test *p*=0.0050. **i-k,** Lack of effects of ICA-121431 on somatic action potential generation. Representative examples of repetitive firing evoked by somatic current injection in sham-control and epileptic mice in ACSF and in the presence of ICA-121431 (blue, panel i). Lack of effects of ICA-121431 on the maximal firing frequency of CA1 neurons (j, k). **l-s,** Effects of the Na_v_1.2/1.6 Na^+^ channel blocker S-Lic (300 μM) on dendritic spikes in sham-control (l-o) and epileptic animals (p-s), depicted in the same manner as in panels (a-h). Two-way ANOVA revealed no significant effects of S-Lic on the rate of rise or threshold of dendritic spikes in control and epileptic animals (m, o). **t-v,** Effects of S-Lic on somatic action potential generation. Representative examples of repetitive firing evoked by somatic current injection in sham-control and epileptic mice in ACSF and in the presence of S-Lic (violet, panel t). Effects of S-Lic on firing induced by current injection at which firing frequency was maximal under ACSF conditions, control and epileptic animals (**u, v,** respectively). Asterisks indicate Bonferroni’s post test *p*=0.0034.

ICA-121431 did not alter the properties of uncaging-evoked single-spine EPSPs (**Fig. S8**). However, in epileptic, but not in sham-control animals, ICA-121431 (100 nM) significantly increased the dendritic spike threshold (**Fig. 3c, g,** spike thresholds indicated with dashed lines, **Fig. 3d, h** for summary, n=5 and 7 in sham-control vs. epileptic, two-way repeated measures ANOVA main effect, sham-control vs. epileptic: F_(1 10)_=0.008, *p*=0.93; ACSF vs. ICA-121431: F_(1 10)_=6.24, *p*=0.032; interaction: F_(1 10)_= 7.10, *p*=0.024; Bonferroni’s post-test epileptic ACSF vs. ICA-121431 *p*=0.0050). Likewise, ICA-121431 decreased the rate of rise of dendritic spikes only in epileptic animals (see insets for dV/dt traces in **Fig. 3a, e**, summary in **Fig. 3b, f,** two-way repeated measures ANOVA main effect, sham-control vs. epileptic: F_(1 10)_=0.050, *p*=0.82; ACSF vs. ICA-121431: F_(1 10)_=5.36, *p*=0.043; interaction: F_(1 10)_=2.50, *p*=0.14; Bonferroni’s post-test epileptic ACSF vs. ICA-121431 *p*=0.025). The NMDA-receptor driven slow phase of the dendritic spike was unaffected by ICA-121431 (two-way repeated measures ANOVA, n.s.).

We then tested if, in addition to the significant effects on dendritic integration in epileptic animals, blocking Na_v_1.3 channels also affects the generation of somatic action potentials. Surprisingly, ICA-121431 had no effects on action potential generation or repetitive firing induced by somatic current injection in either control or epileptic animals (**Fig. 3i-k, Fig. S9, Table S2**, n=6 in both groups, two-way repeated measures ANOVA main effect, sham-control vs. epileptic: F_(1 10)_=11.66, *p*=0.0066; ACSF vs. ICA-121431: F_(1 10)_=1.67, *p*=0.23; interaction: F_(1 10)_=2.28, *p*=0.16, Bonferroni’s post tests n.s.)

We next used the Na^+^ channel blocker and anticonvulsant S-Lic (300 μM), which we found to affect Na_v_1.2 and 1.6 channels, but not Na_v_1.3 or 1.1 channels (*17*) (**Fig. 3l-v**). S-Lic did not affect uEPSPs (**Fig. S8**). Application of S-Lic also did not alter the threshold for eliciting dendritic spikes in either control or epileptic animals (**Fig. 3n, r**, spike thresholds indicated with dashed lines, quantification in **Fig. 3o, s**, n=8 and 5 in sham-control vs. epileptic; two-way repeated measures ANOVA main effect, sham-control vs. epileptic: F_(1 11)_=5.82, *p*=0.044; ACSF vs. S-Lic: F_(1 11)_=0.86, *p*=0.37; interaction: F_(1 11)_= 0.47, *p*=0.51, Bonferroni’s post test n.s.). Likewise, S-Lic did not affect the rate of rise of dendritic spikes in epileptic animals (**Fig. 3m, q,** two-way repeated measures ANOVA main effect, sham-control vs. epileptic: F_(1 11)_=27.76, *p*=0.0003; ACSF vs. S-Lic: F_(1 11)_=1.30, *p*=0.28; interaction: F_(1 11)_= 2.51, *p*=0.14, Bonferroni’s post test n.s.).

As described previously, the Na_v_1.2/1.6 blocker S-Lic significantly inhibited somatic action potential generation (**Fig. 3t-v, Fig. S9**, statistics see Fig. S9 legend). This suggests that the Na_v_1.3 blocker ICA-121431 selectively affects dendritic spikes rather than somatic firing, while S-Lic does the converse.

To elucidate if indeed Na_v_1.3 channels are up-regulated in individual CA1 pyramidal neurons, we investigated *Scn3a* expression at the mRNA level at single-cell resolution. To this end, we used multiplex RNA in-situ hybridization (RNAscope, see Methods, **Fig. S10a,b)**. Low levels of *Scn3a* mRNA puncta were present in pyramidal neurons in the adult hippocampus (**Fig. S10a**). However, we found a significant up-regulation of *Scn3a* expression in pyramidal neurons from epileptic animals (Kolmogorov-Smirnov test p<0.0001 sham-control n=5, epileptic mice n=6, **Fig. S10c**). In contrast, GAD-expressing interneurons showed down-regulation of *Scn3a* compared with sham-control animals (sham-control n=5 epileptic mice n=5, Kolmogorov-Smirnov two-tailed test, p<0.001, **Fig. S10d**).

### Effects of Na_v_1.3 inhibition on altered place-related firing of CA1 neurons

These data point towards a substantial decrease in specificity of dendritic spikes, and degraded input feature detection. We hypothesized that this might contribute to degraded place coding in CA1 neurons in vivo. We therefore examined the activity of CA1 neurons using two-photon Ca^2+^ imaging in head-fixed sham-control and epileptic *Thy1-GCaMP6s* mice running on a linear track equipped with spatial cues (**Fig. 4a-d, Fig. S11a-d**). As described previously, the activity of CA1 neurons was higher during locomotion in both the sham-control and epileptic mice. Additionally, we observed a markedly increased activity of CA1 neurons in epileptic animals compared to sham control animals, both during immobility and locomotion (**Fig. S11e-g,** n=1022 CA1 neurons in 5 sham-control mice and 1697 CA1 neurons in 6 epileptic mice, statistics see figure legend).

**Fig. 4:**
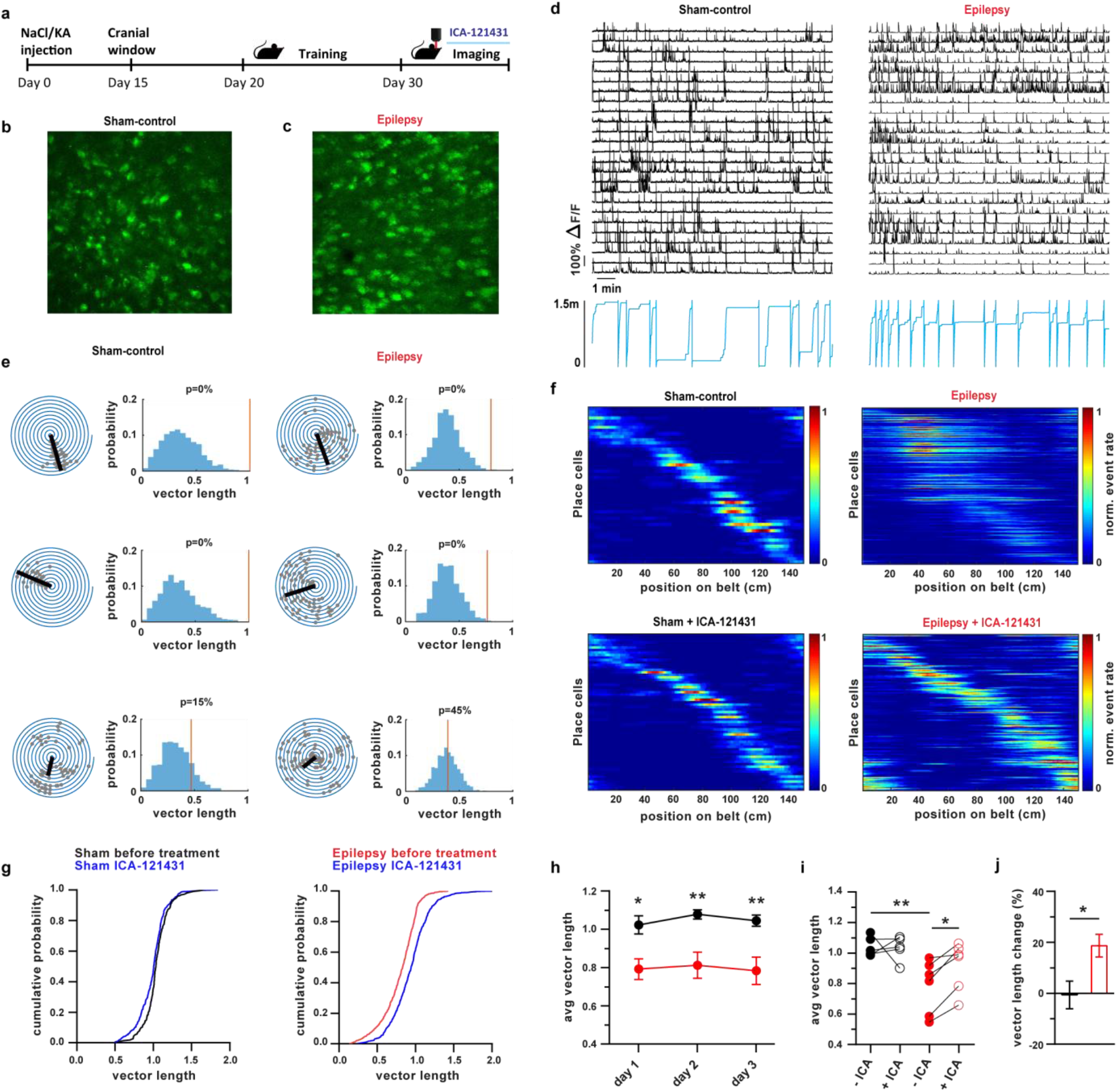
Role of Na_v_1.3 up-regulation in altered place-related firing of CA1 neurons and spatial learning in vivo. **a,** Experimental protocol used to examine the activity of CA1 neurons using two-photon Ca^2+^ imaging in *Thy1-GCaMP6s* mice running on a linear track. **b, c,** Representative fields of view obtained for in vivo imaging. **d,** Representative examples of activity in a subset of the CA1 neurons from the fields of view shown in panels b and c. Upper traces indicate ΔF/F traces from a subset of CA1 neurons, lower blue traces indicate position of the mouse. **e,** Analysis of place coding. Spiral plots for three representative CA1 neurons in a sham-control mouse (left panels) and an epileptic mouse (right panels). One 360° pass around the spiral plot corresponds to a complete transition on the 150 cm circular linear treadmill. Grey circles indicate event onsets in the CA1 neuron. Computed place vectors indicated by black straight lines. Distributions on the right show tests vs. shuffled distributions for each cell (see Methods for details). The red vertical lines indicate the vector length of the CA1 neuron, shuffled distributions shown in blue. **f**, All significantly place-coding CA1 neurons from a representative sham-control (left panels) and epileptic mouse (right panels). Lower panels show the same mice following treatment with ICA-121431. **g,** Cumulative distributions of place vector lengths for sham-control (left panel) and epileptic mice (right panel). Blue curves indicate cumulative distribution of vector lengths after application of ICA-121431. **h,** Differences between average vector lengths in sham-control (black) and epileptic animals (red) were stable over imaging sessions on three consecutive days. Two-way ANOVA main effect, sham-control vs. epileptic: F_(1 27)_=31.12, p<0.0001; subsequent days: F_(2 27)_=0.2623, *p*=0.7727; interaction: F_(2 27)_=0.05973, *p*=0.9421; asterisks indicate Bonferroni’s post-test, day 1 *p*=0.02, day 2 *p*=0.0066 and day 3 *p*=0.0074. **i,** Average vector lengths calculated per mouse in sham-control (black) and epileptic mice (red). Asterisks indicate Bonferroni’s post-tests sham-control vs. epileptic *p*=0.0095, epileptic mice before ICA-121431 treatment vs. ICA-121431 treated epileptic mice *p*=0.0207. **j,** Percent change in vector length caused by ICA-121431, quantified for each animal and averaged. Asterisk indicates Mann-Whitney U-test, *p*=0.030.

CA1 neurons that exhibited significant place-related activity (*18*) were found in both sham-control and epileptic mice (examples of representative neurons in **Fig. 4e,** spiral plots show place coding, rightmost distributions show tests vs. shuffled distributions for each cell, more examples of individual neurons in **Fig. S12**). In both sham-control and epileptic mice, place-related firing fields tiled the extent of the linear track (**Fig. 4f**, upper panels all significantly place-coding CA1 neurons from a representative sham-control and epileptic mouse, respectively**, Fig. S13** for data from all sham-control and epileptic mice). It was apparent from visual inspection of these data that place-related firing of significantly place-coding cells appeared to be less specific in epileptic animals, consistent with published work demonstrating degraded place coding in epileptic (*19–22*). We quantified the precision of spatial coding using a spatial tuning vector measure ((*18, 23*), see also place vectors in spiral plots in **Fig. 4e**), where higher place coding specificity corresponds to greater vector lengths. Indeed, we found the distribution of place vector lengths was significantly shifted to shorter values in epileptic mice, implying degraded place coding (**Fig. 4g,** cf. left and right panel). This difference was stable over multiple sessions of imaging (**Fig. 4h**), indicating decreased specificity of place coding in epileptic animals. In the presence of ICA-121431, vector lengths significantly increased in epileptic, but not in sham-control mice (cumulative distributions of vector lengths for all cells in **Fig. 4g**). Correspondingly, ICA-121431 also caused an increase of the average vector lengths calculated per mouse in epileptic, but not in sham-control mice (**Fig. 4i,** n=6 and 5 mice respectively, repeated-measures two-way ANOVA main effects, sham-control vs. epileptic: F_(1 9)_=6.29, *p*=0.033; baseline vs. ICA-121431 treatment: F_(1 9)_=3.92, *p*=0.079; interaction: F_(1 9)_=5.64, *p*=0.042; Bonferroni’s post-test sham-control vs. epileptic *p*=0.0095, epileptic mice before ICA-121431 treatment vs. ICA-121431 treated epileptic mice *p*=0.0207). When we computed the effects on average vector lengths in each animal, the effect of ICA-121431 was significantly larger in epileptic mice, as expected (**Fig. 4j,** Mann-Whitney U-test, *p*=0.030). These experiments suggest that blocking Na_v_1.3 channels in vivo significantly reverses degraded place coding in epilepsy.

It is known that place coding can be disrupted by interictal synchronous activity (*24, 25*). To elucidate if effects of ICA-121431 on place-related firing in epileptic animals could also be due to an inhibition of aberrant synchronous activity, we tested the effects of this compound on population activity in the CA1 region. We found aberrant synchronized activity in epileptic mice, but never in control mice (Event frequency in epileptic mice 0.64±0.074/minute, example in an epileptic mouse shown in **Fig. S14a**). Neither the number of aberrant events (**Fig. S14b-d**) nor the duration of the events (**Fig. S14c, e**) was affected by ICA-121431 (see Supplementary Materials for analysis methods, Fig. S14 legend for statistics).

### *Inhibiting Nav1.3 channels normalizes hippocampal-dependent memory in* epileptic mice

We next asked if inhibition of Na_v_1.3 channels with ICA-121431 in vivo restores performance in a hippocampus-dependent spatial memory task. We selected a task in which rodents learn the spatial arrangement of two objects. In a subsequent session, one of the objects is displaced (object location memory test, OLM, **Fig. 5**), and the increased exploration of the displaced object is measured. In subsequent experiments, one of the objects is exchanged for a novel object (novel object recognition, NOR test, see **Fig. S15**). After habituation for three days, treatment was started with either vehicle or ICA-121431 during the fourth habituation day (**Fig. S15b**). Subsequent sessions for the OLM test were carried out either in the presence of vehicle or ICA-121431 (**Fig. 5a**). During acquisition of the object position for the OLM test, mice did not discriminate the two objects (**Fig. 5b, c,** upper panels, **Fig. 5d,** two-way ANOVA, n.s.). In the OLM trial with a displaced object, vehicle-treated sham-control mice discriminated the displaced object, while vehicle-treated epileptic animals did not (**Fig. 5b,** lower panels, n=17 and 15, respectively, filled bars in **Fig. 5e**). Importantly, administration of ICA-121431 by gavage (see methods) recovered memory performance to control levels in epileptic animals (**Fig. 5c,** lower panels, n=7 and 8 for sham-control and epileptic animals, respectively, empty bars in **Fig. 5e**). Two-way ANOVA revealed a main effect of vehicle vs. ICA-121431 treatment (vehicle vs. ICA-121431: F_(1 43)_=5.95, *p*=0.019; sham-control vs. epileptic: F_(1 43)_=1.17, *p*=0.29; interaction: F_(1 43)_=3.91, *p*=0.054 Bonferroni’s post tests indicated with asterisks).

**Fig. 5.**
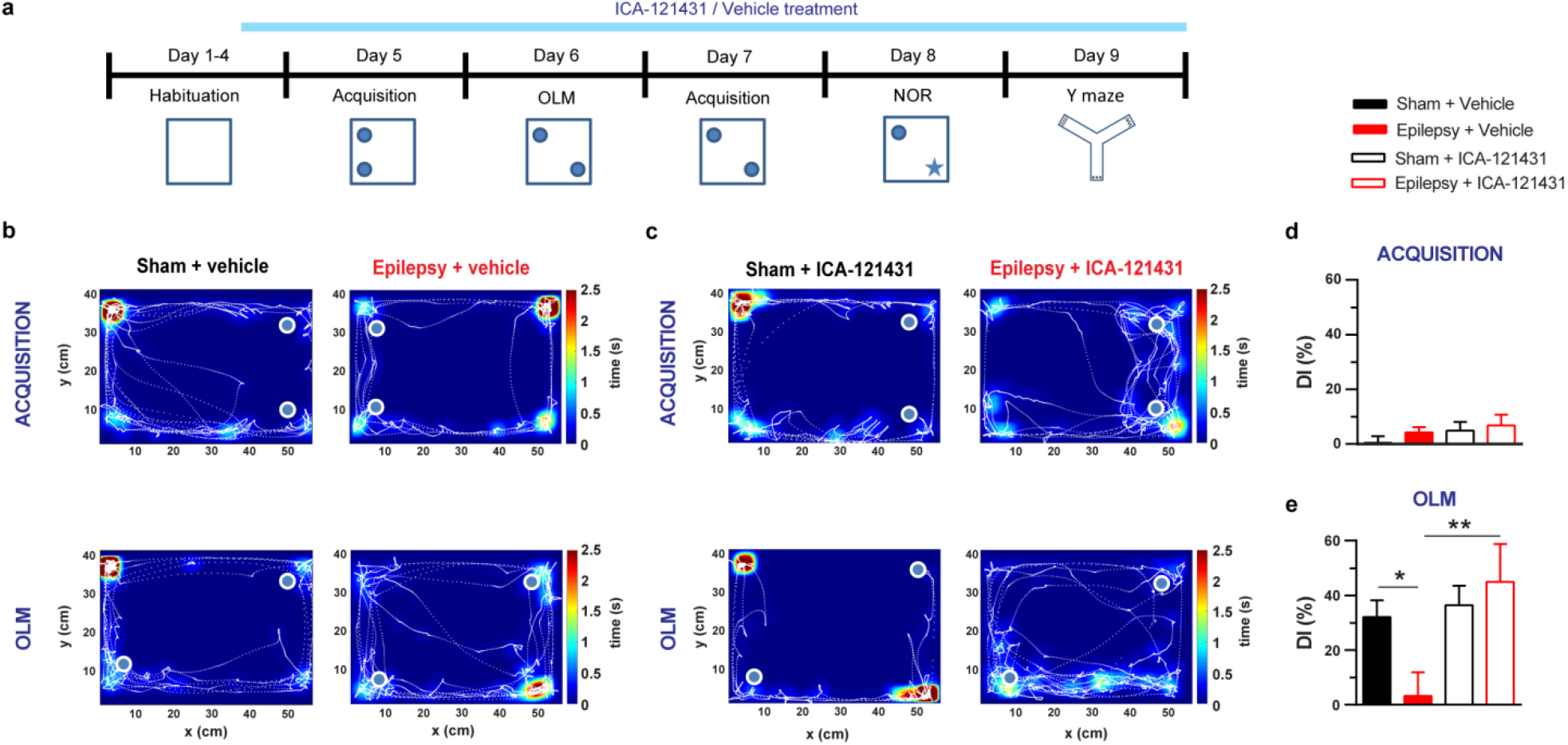
Inhibiting Na_v_1.3 channels normalizes hippocampal-dependent memory in epileptic mice. **a,** Timeline of the behavioral experiments with in vivo inhibition of Na_v_1.3 channels with ICA-121431. After habituation to the open field, the object location memory (OLM), object recognition memory (NOR) test and Y maze spontaneous alternation test were sequentially performed (**Fig. S15)**. **b, c,** Representative examples of sessions with tracking data in sham-control and epileptic animals with vehicle application (b) and ICA-121431 application (c), respectively. Occupancy times are color coded (calibration see scale bar). **d,** The discrimination index (DI) indicating if mice preferentially explore an object during the acquisition session. DI was close to zero, indicating that mice did not discriminate the two objects in any group. **e,** DI in the OLM session. Vehicle-treated sham-control mice discriminated the displaced object, while vehicle-treated epileptic animals did not (black and red filled bars, respectively). Administration of ICA-121431 by gavage recovered memory performance to control levels in epileptic animals (empty bars). Asterisks indicate Bonferroni’s post-test, sham-control vehicle treated vs. epileptic vehicle treated *p*=0.0193; epileptic vehicle treated vs. epileptic ICA-121431 treated *p*=0.0057.

In contrast to the OLM trial, novel object recognition (NOR) was not impaired in epileptic animals (**Fig. S15c, d**), consistent with previous reports (*12*). ICA-121431 treatment did not alter performance in the NOR trial in either sham-control or epileptic mice (**Fig. S15e, f,** n-numbers as for OLM trials, statistics see Fig. S15 legend).

We next investigated if pharmacological inhibition of Na_v_1.3 channels with ICA-121431 affects performance in a spatial working memory task, spontaneous alternation in the Y-maze. In this task, animals freely explore a three-arm maze. The extent to which animals sequentially explore three different arms is quantified (alternation score), with higher alternation scores indicating a higher level of efficient exploration requiring spatial working memory (**Fig. S15i**). ANOVA revealed main effects on the alternation score of kainate treatment (sham-control vs. epileptic: F_(1 44)_=7.36, *p*=0.0095), but the effects of ICA-121431 treatment did not reach statistical significance (F_(1 44)_=2.86, *p*=0.098; interaction: F_(1 44)_=0.82, *p*=0.59, **Fig. S15k**, Bonferroni’s post tests indicated with asterisks). Thus, the effects of ICA-121431 seem to be most prevalent for spatial tasks that require hippocampal-dependent memory consolidation.

## Discussion

In this paper, we describe a Na^+^ channel dependent mechanism underlying a prominent change in dendritic supralinear integration in epilepsy that degrades place coding in vivo, and causes deficits in spatial memory.

Dendritic spikes are a cornerstone of dendritic integration that enable neurons to generate precise action potential outputs to clustered inputs deriving from specific populations of presynaptic input neurons. They are therefore important in mediating sharply tuned neuronal responses to features of input neurons (*1, 26*). We show that the properties of dendritic spike generation are markedly altered in CA1 pyramidal cells in chronic epilepsy. Specifically, the fraction of dendritic branches that can give rise to dendritic spikes is almost doubled in epileptic animals. In addition, in those dendrites that generate dendritic spikes, virtually all properties that confer input-specificity to dendritic spikes are degraded. First, even very asynchronous stimulations can cause dendritic spikes in epileptic animals. Second, the threshold for eliciting dendritic spikes is significantly lowered in epileptic animals. Third, mechanisms that usually attenuate dendritic spikes upon prior occurrence of somatic or dendritic spikes (*14*) are much less effective in epileptic animals. Fourth, dendritic spikes rise at an enhanced rate in 1^st^ order dendrites from epileptic animals. Because dendritic spikes with a high rate of rise (also termed ‘strong’ spikes) are much less affected by concurrently evoked inhibition (*27*), this predicts decreased inhibitory control of dendritic spikes in epileptic animals. The changes in dendritic integrative properties were primarily present in 1^st^ order, and not 2^nd^ order dendrites. This is reminiscent of the changes observed following environmental enrichment, in which dendritic spikes in 1^st^ order branches are selectively strengthened (*28*). In this paradigm, strengthening of proximal branches led to an enhanced propagation of distally evoked dendritic spikes into 1^st^ order dendrites (*28*). Importantly, epilepsy-related changes in dendritic integration are observed in CA1, but not in other hippocampal subregions such as the dentate gyrus (*29*).

Together, these findings suggest that input feature detection of CA1 neurons is degraded in epileptic animals, and predict much less selective task- or stimulus-specific response properties. We have used place coding as a model to test this idea. Place cells within the CA1 fire specifically when a rat occupies a particular location in the environment (*30*). Dendritic spiking has been proposed to be essential in driving formation of place-related firing in CA1 neurons (*4, 31, 32*), and computational studies have suggested that sharp spatial tuning of place cell responses is supported by dendritic spikes (*33*). Indeed, we have found significant broadening of place representations in epileptic animals, consistent with previous studies in different models of epilepsy (*19–21, 34*). This idea is also in line with computational studies suggesting that changes causing an increase in dendritic spike generation degrade place tuning of pyramidal neurons (*33, 35*). Importantly, the absence of dendritic spikes also degrades spatial tuning, suggesting that the prevalence and properties of dendritic spikes have to be carefully tuned for optimal place coding (*35*).

Our pharmacological data show that enhanced excitability of 1^st^ order dendrites is most likely caused by up-regulation of Na_v_1.3 channels. This channel isoform exhibits rapid repriming and recovery from inactivation, as well as particularly slow closed-state inactivation, properties that are well suited to explain repetitive generation of dendritic spikes, and loss of dendritic spike inactivation (*36*). The up-regulation of these channels is consistent with a proximal dendritic up-regulation of persistent Na^+^ channels described in a different model of experimental TLE (*10*). Although the convergent in vitro and in vivo data point to a role of Na_v_1.3 channels, additional changes also influence processing of CA1 neurons during navigation. Inhibition is strongly altered in chronic epilepsy, with changes in the amount and timing of inhibition delivered by inhibitory circuits (*37–39*), a factor that may also contribute to altered place coding. However, it has been argued that the development of interneuron loss and seizures occurs long before instability of place representations is observed (*21*), suggesting that interneuron loss alone cannot account for this phenomenon. It is nonetheless likely that changes in inhibitory circuits act together with degraded specificity of dendritic spikes to compromise the precision of CA1 firing.

Could the up-regulation of Na_v_1.3 be therapeutically targeted to treat cognitive comorbidities of chronic epilepsy? Several avenues have been shown to be potentially feasible. Despite the high degree of structural similarity between individual Na^+^ channel isoforms, selective antagonists for specific Na^+^ channel subunits have been developed, including for Na_v_1.3 (*16*). In addition, antisense oligonucleotides could be used to selectively inhibit Nav1.3 expression, potentially in a cell-type specific manner (*40, 41*). The availability of anticonvulsants that inhibit other, non-Na_v_1.3 Na^+^ channel isoforms, such as S-Lic, and have a proven efficacy against partial seizures, may also allow to combine blockers affecting abnormal dendritic spiking vs. somatic action potential firing. In addition to applying direct Na_v_1.3 modulators, the biological mechanisms leading to enhanced Na_v_1.3 expression could also be targeted. A potential mechanism for the up-regulation of Na_v_1.3 in epileptic animals is a glycerinaldehyd-3-phosphate-dehydrogenase (GAPDH) mediated posttranscriptional regulation (*15*). Importantly, this up-regulation is ameliorated by ketogenic diet, which is used to control therapy-refractory epilepsies in children (*15*). In addition, the anticonvulsant valproic acid has been shown to epigenetically reduce Na_v_1.3 expression via promoter methylation (*42*). Thus, several avenues for direct modulation of Na_v_1.3 channels or regulation of their expression levels may be exploited to correct epilepsy-induced Na_v_1.3 overexpression, and to reverse cognitive impairments in chronic . TLE.

## Acknowledgements

We are very grateful to David Greenberg, Jason Kerr, and Damian Wallace for technical help and advice, as well as the supply of analysis algorithms. We gratefully acknowledge the support of Jonathan Ewell in editing the manuscript. We are grateful to Lea Keller for technical support, and Thoralf Opitz for support with animal husbandry and in-vivo pharmacology. We acknowledge the support of the Imaging Core Facility of the Bonn Technology Campus Life Sciences funded by the Deutsche Forschungsgemeinschaft (DFG, German Research Foundation) – Projektnummer 388169927.

## Funding

The work was supported by the SFB 1089 to HB, SB and CH, the Research Group FOR2715 to TK and HB, and SPP 2041 to HB, and EXC 2151 under Germanys Excellence Strategy of the Deutsche Forschungsgemeinschaft (DFG, German Research Foundation) to HB, BL 767/5-1 to SB and support of the Humboldt Foundation PSI to KG. ANH was supported by the IMPRS Brain and Behaviour.

## Author contributions

H.B., N.M., and T.K. conceived and planned the study. N.M. performed electrophysiological, uncaging and in-vivo imaging experiments and data analysis, N.N. and M.P. designed the linear treadmill, head fixation and hippocampal window, and contributed to the data analysis. K.B. and C.H. did electrophysiological and in-vitro imaging experiments and contributed to experimental design. M.P. and L.E. did in-vivo electrophysiological experiments. F.K. did analysis of calcium imaging data. K.U.S.I. and S.B. contributed RNAscope experiments. A.N.H. and K.G. performed and analyzed behavioral experiments and designed the behavioral apparatus, N.M., T.K., and H.B. wrote the manuscript. All authors participated in the preparation of the manuscript.

## Competing Interests

The authors declare no competing interests. N.M. was partially financed by BIAL.

## Data availability

All data in the paper will be made available upon reasonable request.

## Supplementary Materials

Materials and Methods

Figures S1-S15

Tables S1, S2

## Notes

### Competing Interest Statement

The authors have declared no competing interest.

### Summary of Updates

Author affiliations updated Supplemental files updated

